# When are Biomedical Postdocs Ready for the Faculty Job Market? A Mixed-Methods Analysis of Metrics and Resilience Among Faculty Job Seekers

**DOI:** 10.64898/2026.08.14.744880

**Authors:** Amanda Haage, You Cheng, Christopher T. Smith, Ariangela J. Kozik, Ada K. Hagan, Nafisa M. Jadavji

**Author notes:** Co-corresponding authors, Corresponding Authors: Nafisa M. Jadavji, Division of Molecular and Integrative PhysiologyDepartment of Biomedical Sciences, School of Medicine, Southern Illinois University, Mail Code 6512 1135 Lincoln Drive, Carbondale, IL 62901, Amanda Haage, Department of Biomedical Sciences, University of North Dakota School of Medicine and Health Sciences, 1301 N Columbia Rd Grand Forks, ND 58202 Stop 9037, You Cheng, McLean Imaging Center, McLean Hospital 115 Mill St. Belmont, MA 02478.

## Abstract

**Purpose:** Discussions surrounding the biomedical faculty job market often focus on applicant competitiveness and external metrics such as number of publications and funding records. Consequently, there is typically less discussion about applicant readiness, the point at which applicants perceive themselves as prepared to enter the market. Since 2018 our group, the Faculty Job Market Collaboration (FJMC), has conducted annual end-of-cycle surveys of biomedical faculty job applicants, producing the largest longitudinal dataset on this process to date.

**Methods:** We employed a mixed-methods design examining faculty applicants in biological science fields in North America. Regression analyses were conducted on a longitudinal dataset of 729 respondents across multiple hiring cycles. To determine how applicants evaluated their own preparation, qualitative interviews were conducted with a separate cohort of biomedical postdoctoral applicants during the 2024-2026 job cycles.

**Results:** Our findings demonstrate that rather than depending on a single quantitative threshold, readiness is a multifaceted construct shaped by actionable and interpersonal drivers. Key factors influencing an applicant’s perceived readiness include taking agency to submit applications, receiving explicit support from a mentor, incorporating strategic use of artificial intelligence tools into application preparation, and their career stage.

**Conclusion:** By distinguishing individual readiness from systemic assumptions of market competitiveness, this study highlights a blind spot in academic workforce development. Our results suggest that applicants can achieve readiness and successful outcomes through different combinations of support, strategy, and timing rather than a uniform metric profile. By integrating quantitative and qualitative data, our study provides an evidence-based framework for understanding applicant readiness and offers practical guidance to help trainees navigate the increasingly competitive academic job market.

**Teaser Text:** Our mixed-model analysis of the biomedical faculty job market is designed to help prospective faculty candidates assess their readiness to enter the job market. By integrating multiple indicators of academic productivity, funding success, and professional experience, our study provides evidence-based benchmarks that can guide applicants in evaluating their competitiveness and identifying areas for further development before pursuing faculty positions.

## Introduction

The transition from postdoctoral training to an independent faculty position represents one of the most consequential career decisions in academic biomedicine, shaping the future research, teaching, and mentoring workforce. Despite its importance, the faculty job market has historically been characterized by limited empirical evidence and heavy reliance on anecdotal guidance. Over the past several years, the Faculty Job Market Collaboration (FJMC) has helped address this gap through a series of national studies describing applicant experiences, identifying predictors of hiring outcomes, and examining trends in academic hiring^1–4^. Collectively, this work demonstrates that no single metric defines a successful applicant, and while scholarly productivity, funding, and application behaviors are all associated with faculty hiring outcomes, evaluating faculty job market competitiveness is complex.

Yet identifying predictors of hiring success does not explain how applicants decide they are ready to enter the faculty job market. Career development theories suggest that readiness extends beyond accumulated accomplishments to include an individual’s preparedness to make informed career decisions within complex and uncertain environments^5^. From this perspective, readiness emerges through individuals’ interpretation of their accomplishments, perceived capabilities, and the demands of the career transition they are preparing to undertake. Importantly, competitiveness and readiness represent related but distinct constructs: competitiveness reflects the qualifications and experiences associated with success in the faculty job market, whereas readiness reflects an applicant’s judgment that they are prepared to enter it^6^. Within the biomedical sciences, Clement and colleagues (2020)^7^ advanced this concept through the Academic Career Readiness Assessment (ACRA), an evidence-based framework that defines readiness from the perspective of faculty hiring committees by identifying the qualifications and levels of achievement expected across institution types. Although ACRA provides an important foundation for mentoring and faculty career preparation, it conceptualizes readiness in terms of institutional expectations rather than applicants’ decisions about when to enter the faculty job market. Consequently, little empirical evidence exists to explain how applicants integrate objective indicators of competitiveness with their own perceptions of preparedness when deciding if they are ready to apply.

The purpose of this study was to examine readiness to enter the faculty job market from the applicant perspective using a sequential explanatory mixed-methods design. We first analyzed national survey data from biomedical faculty job applicants to identify applicant characteristics that predicted obtaining a faculty job offer. We then integrated findings from the first interview in an ongoing four-interview longitudinal study, capturing applicants’ reflections as they entered the faculty job market. Interview data were analyzed using a structured coding framework to characterize perceived preparedness across key domains of faculty career development, including research, funding, teaching, mentoring, networking, mentorship, interdisciplinary training, and personal context. By integrating quantitative and qualitative findings, this study extends prior work on predictors of hiring success to develop an empirically grounded framework for understanding how applicants evaluate their readiness to pursue faculty careers.

## Methods

### Study Design

This study employed a convergent mixed-methods design, combining longitudinal survey data from six hiring cycles (2019 to 2025) with semi-structured interview analysis conducted prospectively during the 2024-25 and 2025-26 faculty job market cycles. Survey (quantitative) and interview (qualitative) components were analyzed separately and synthesized at the interpretation stage; an analysis pipeline has been outlined in **Figure 1**.

**Figure 1.**
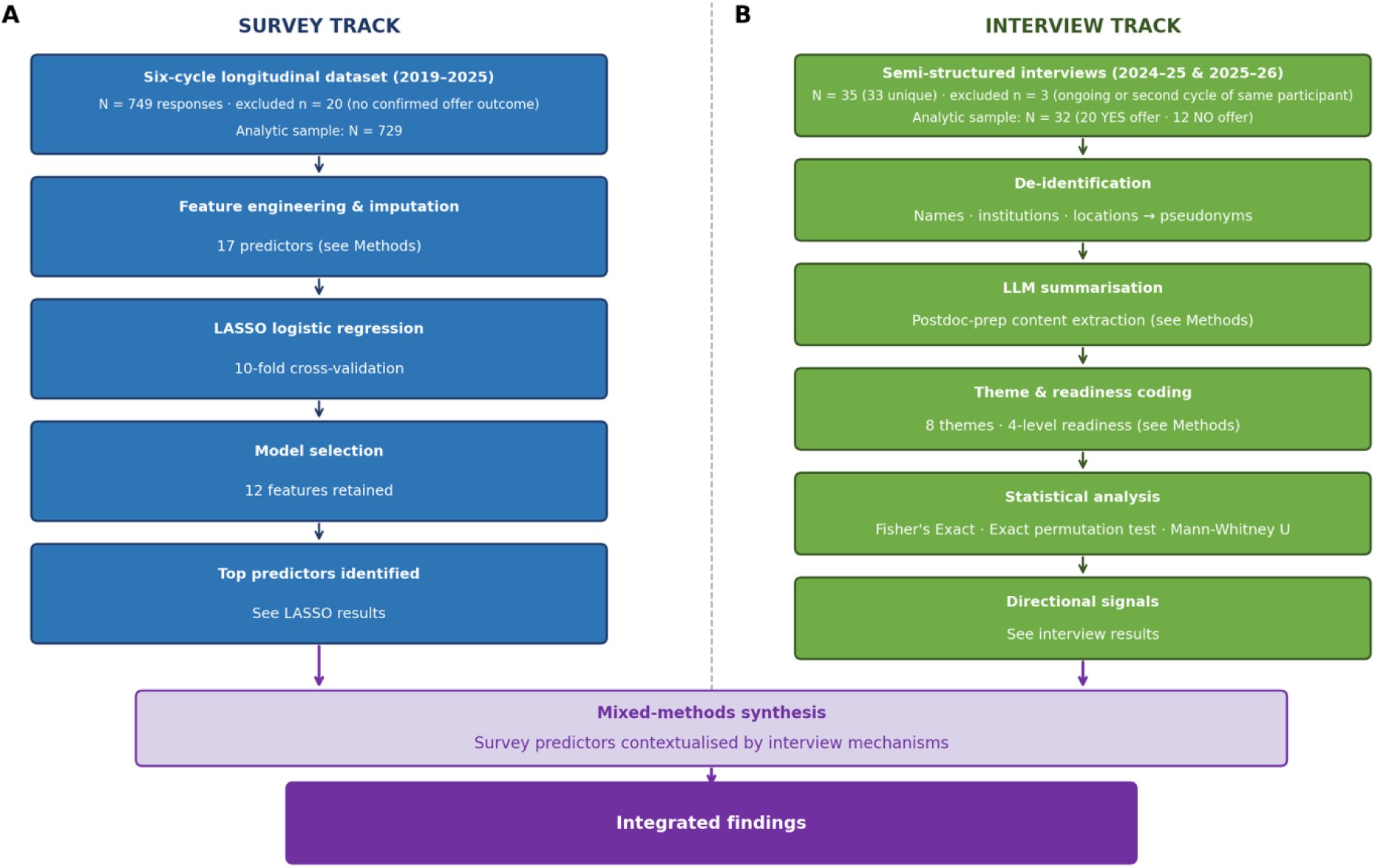
Data analysis pipeline overview completed for mixed-methods synthesis of biomedical faculty job market data. The survey track arm (A) gives an overview of the LASSO workflow. Survey arm (n = 729) responses with confirmed offer outcome across 6 hiring cycles, 2019 –20; 2020-21; 2021-22; 2022-23; 2023-24; 2024-25. The interview track arm (B) gives an overview of large language models (LLM) + coding + statistical analysis pipeline. Both tracks converge in mixed-methods synthesis. Interview arm (n = 35) transcripts from interview one during 2024-25 and 2025-26 job cycles. Outcomes of all included participants were known, received faculty job market offer or did not. Three participants were excluded because one was still participating in faculty job market interview processes at the time of data analysis and one participant was interviewed twice.

The quantitative study was approved by the University of North Dakota Institutional Review Board (IRB project number: IRB-201908-045) as exempt according to 4 5CFR46.101(b)(2): anonymous surveys no risk on 08/29/2019. The qualitative interview study was approved by Southern Illinois University (IRB project number: 24-587) as exempt under 45 CFR 46.104(d) and 38 CFR16.104(b) on 10/02/2024, University of Michigan (IRB project number: IRB00000246) as exempt according to 2(i) and/or 2(ii) at 45 CFR 46.104(d) on 09/16/2024 and Virginia Polytechnic Institute and State University (IRB project number: 24-889) as exempt according to 45 CFR 46.104(d) category 2(ii) on 12/09/2024.

### Survey Data Collection

Faculty applicants in the biological sciences in North America completed a self-report survey during six consecutive hiring cycles spanning academic years May 2019 through November 2025 (**see supplementary file for survey questions**). Data from the initial three cycles (2019-2020; 2020-2021; 2021-2022) were collected and reported in earlier work from our group^1^. The present study adds three new cycles (2022-2023; 2023-2024; 2024-2025), yielding a combined analytic sample of n = 729 (per-cycle n = 231 (2019-2020), 64 (2020-21), 154 (2021-22), 97 (2022-23), 76 (2023-24), 107 (2024-25). The survey was distributed via email to postdoctoral administrator offices and organizations, as well as through social media platforms including Bluesky, LinkedIn, and Twitter (X). Respondents were excluded if they did not meet a 33% minimum completion threshold, reported prior tenure-track employment, or did not disclose their faculty offer outcome.

Survey items assessed: self-reported demographics (gender identity [coded 0 = Man; 1 = Woman/LGB+/GNC (trans/gender non-confirming)], age category, international residency, disability status, first-generation undergraduate and doctoral status); career-stage characteristics (current academic position, number of prior application cycles, total applications submitted per cycle, number of postdoctoral positions held); and publication metrics (number of first-author papers, total peer-reviewed publications, h-index, total citations). Peer mentorship status was also assessed.

### Interview Participants

Semi-structured interviews were conducted with faculty applicants during the 2024–2025 and 2025–2026 hiring cycles by trained members of the Faculty Job Market Collaboration (FJMC) team (**Figure 1B**). Applicants completed a minimum of three semi-structured interviews focused on job market preparation, virtual interview readiness, and on-site interview preparation. Candidates who did not progress beyond specific stages of the search process participated in a final interview centered on future career plans. Those who received faculty offers completed an additional interview addressing offer negotiation and informed decision-making. Questions from the first interview (supplement) which included preparation for the faculty job market were used for this study and outcomes of whether an applicant received a job offer or not were included in our analysis (**see supplementary information for questions**).

Transcripts were de-identified prior to analysis, replacing all proper nouns (participant names) with pseudonyms or neutral descriptors. Institution names were labelled as classified by the Carnegie classification. Geographic identifiers were replaced with a broad US region. Participants were matched to confirmed faculty offer outcomes using the FJMC applicant tracking database. Inclusion required both a complete de-identified transcript and a confirmed offer outcome. Applicants with an ongoing application cycle at the time of analysis (n = 1) and second-cycle transcripts for participants whose first-cycle interview was already included (n = 2) were excluded to avoid double-counting. The final analytic interview sample comprised n = 32 participants: 20 with a confirmed faculty offer (YES) and 12 without (NO), yielding an observed offer rate of 62.5%. The participant numbers are outlined in **Figure 1B**.

### Interview Coding

Transcripts were processed through a standardized four-step pipeline: verbatim transcription, chunked LLM-assisted summarization to extract postdoctoral preparation narratives (Mistral 7B via Ollama 0.5.3, executed locally on an Apple M1 Max MacBook Pro [10-core CPU, 32-core GPU, 64 GB unified memory] with Metal GPU acceleration; no transcript data were transmitted to external servers), regex-based theme detection, and LLM-assisted readiness classification. Eight thematic domains were coded: Research Productivity, Grantsmanship, Teaching/Mentoring, Networking, Cohort Support, Personal Context (Family/Geography), Mentorship (PI), and Research Interdisciplinarity. Readiness in each coded theme was assigned on a four-level scale: Prepared (narrative indicated adequate preparation), Mixed (partial preparation with co-occurring gaps), Unprepared (clear preparation deficits identified), or Unknown (insufficient transcript content to assign a rating). Each participant received one readiness code per identified theme.

### Statistical Analysis

For survey data, faculty offer receipt (0 = no offer; 1 = at least one offer) was modeled using LASSO logistic regression (L1 penalty, liblinear solver) and feature selection based on previous work by our group (Flynn et al). Missing predictor values were imputed using k-nearest-neighbor imputation (k = 5) prior to standardization. The postdoc count item showed markedly increased non-response in cycles 4–6 (22%–29%) relative to cycles 1–3 (<8%) because it was marked as optional in cycles 4 to 6; to test whether non-response itself carried a predictive signal. The regularization parameter C was selected by 10-fold cross-validation over a grid (C ∈ [0.10, 0.32], step 0.01), optimizing mean cross-validated area under the receiver operating characteristic curve (AUC). Cycle-level model stability was assessed by fitting the same LASSO pipeline independently within each hiring cycle (**Supplementary Figure 1**). Analyses were conducted in Python 3.10.8 (scikit-learn 1.4.2, numpy 1.25.2).

Interview data were analyzed using descriptive and inferential methods. Readiness distributions across the eight thematic domains were examined descriptively; participants coded Unknown for a given theme were excluded from theme-level analyses. The proportion of participants with at least one theme coded as Prepared was compared between offer groups descriptively, as the small analytic sample precluded formal statistical testing. Participants were asked about AI usage and 25 out of the 32 responded. The distribution of use categories (none, writing/editing, research/search, multiple uses) was compared between YES-and NO-offer groups using Fisher’s exact test (two-tailed). Analyses were conducted in Python 3.10.8 (SciPy 1.9.3, pandas 2.3.3).

## Results

### Predictors of a faculty job offer

Since 2019 we have sought to bring transparency to the faculty job market by using yearly quantitative surveys describing diverse applicant metrics and their outcomes. This has allowed us to maintain a public dashboard of this data^8^ and perform LASSO logistic regression analyses^3^. Here we present our most up-to-date LASSO logistic regression combining six years of applicant data (**Figure 2**). The performance of our LASSO regression had lower performance in cycles with smaller numbers of respondents (**Supplementary Figure 1**). We have shown repeatedly, in 2018^2^ and 2022^3^, that the number of applications per cycle (β = +0.67) impacts whether applicants receive a job offer. Additionally, a more senior position is favourable (β = +0.61), such as a research scientist or research assistant professor, as well as other senior position. When an applicant has completed many application cycles (β = −0.12), is older (β = −0.14), and has held more postdoc positions (β = −0.15), these attributes are negatively associated with receiving a job offer. Knowledge of the academic system and the hidden curriculum^9^ also appears to be beneficial, as our data show that being a first generation undergraduate (β = −0.05) or PhD (β = −0.02) and having an international residence (β = −0.08) can reduce the odds of receiving a job offer. Together, this data paints a picture where there is a specific window when applicants are most competitive: senior but not held too many post-PhD positions and that they need to apply broadly, as higher application numbers are associated with greater success in receiving an offer in our data. Our survey data suggest that applicants with more inherent knowledge of academic systems may be able to identify better when that window occurs in their career. This led us to ask how do applicants decide when they are ready to enter the faculty job market?

**Figure 2.**
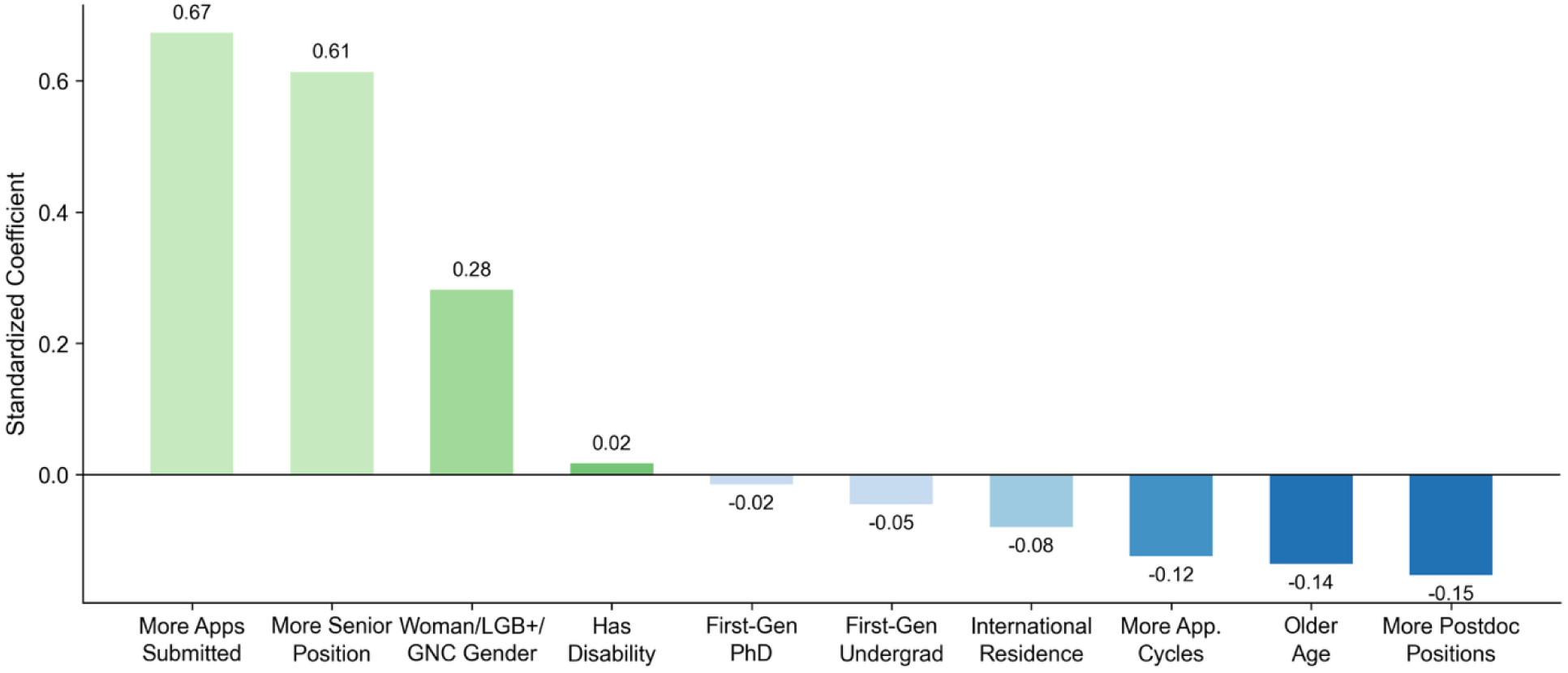
Regression model analysis (LASSO standardized coefficients) showing the relationship between factors and receiving faculty job offers in the biological sciences over five cycles (2019 –20; 2020-21; 2021-22; 2022-23; 2023-24; 2024-25) with a sample size, n = 729). Positive coefficients (light green) indicate association with receiving a faculty offer and negative coefficients (blue) indicate negative association. We have identified an optimal regularization parameter of C = 0.2, yielding a mean cross-validated AUC of 0.73 ± 0.04. Of 16 candidate features, 10 were retained with non-zero coefficients. The three strongest positive predictors were application volume (β = +0.67), career seniority (β = +0.61), and non-Man gender identity (β = +0.28). Negative predictors included number of postdoctoral positions held (β = −0.15), age (β = −0.14), number of prior application cycles (β = −0.12), international residency (β = −0.08), first-generation undergraduate status (β = −0.05), and first-generation doctoral status (β = −0.02). Disability status was retained at a small positive coefficient (β = +0.02). Total peer-reviewed publications, h-index, citation count, first-author publication count, peer mentorship, and number of dependents were not selected (β = 0).

### Factors that influence applicant perception of preparedness

Participants for our qualitative study were drawn from the 2024–2025 and 2025–2026 hiring cycles. A total of 32 interview participants were recruited, 20 received at least one faculty offer (62.5% offer rate). Using transcripts from interview one, eight thematic domains were derived from the topic areas addressed systematically across the semi-structured interview protocol: research productivity, grantsmanship, teaching/mentoring, networking, cohort support, personal context (family/geography), mentorship by PI, and interdisciplinarity. These domains reflect constructs the protocol was designed to elicit at each stage of the faculty job search, enabling direct comparison across participants.

We coded interview responses for all 32 analytic participants across eight faculty preparation themes (**Figure 3A**). Mentorship by applicant’s PI was the most frequently coded theme (n = 32). It showed the highest combined proportion rated by applicants feeling prepared (31%) or mixed, neither prepared nor unprepared (41%) about the faculty job market. While networking showed a more even distribution across perceived readiness levels.

**Figure 3.**
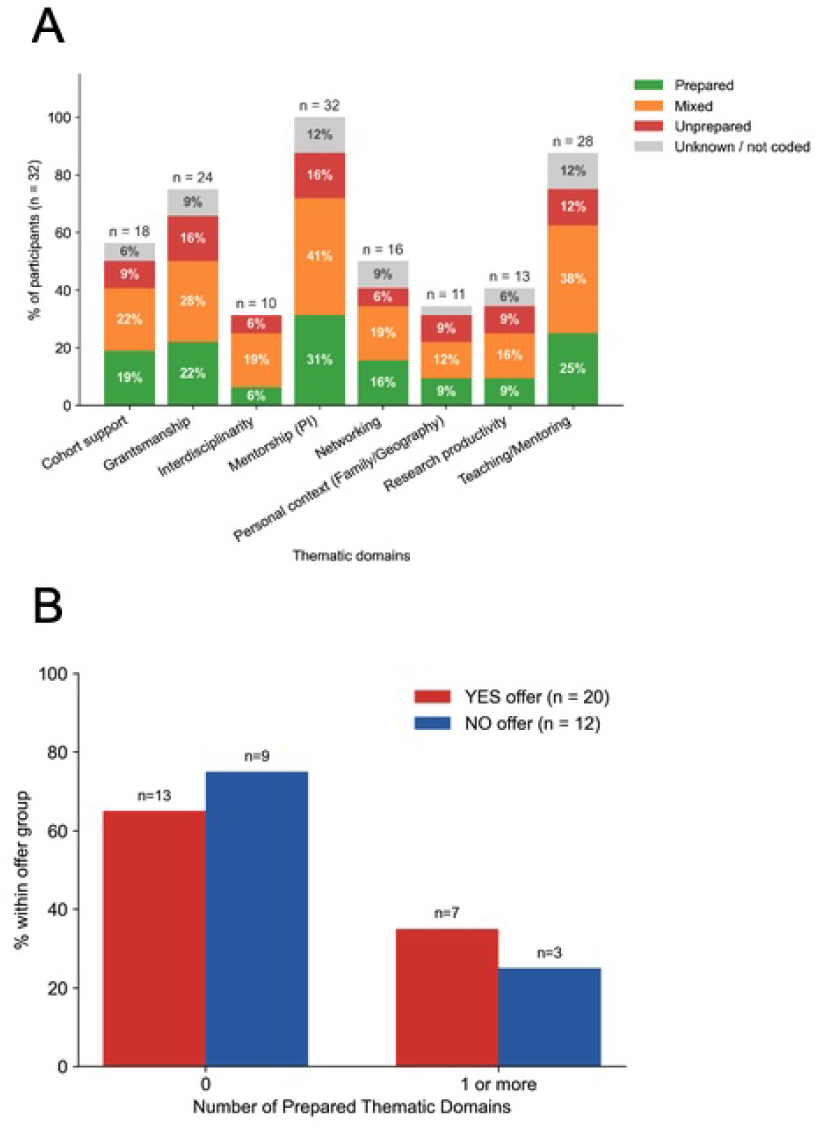
Interview one analysis of biomedical sciences postdoctoral fellows on faculty job market (participants). Readiness distribution (A) by theme across all 32 interview participants. Bar height reflects the percentage of all analytic participants (n = 32). Eight thematic domains were coded: Research Productivity, Grantsmanship, Teaching/Mentoring, Networking, Cohort Support, Personal Context (Family/Geography), Mentorship (PI), and Interdisciplinarity. Readiness in each coded theme was assigned on a four-level scale: prepared (green), mixed (orange), unprepared (red), or Unknown (grey). Each participant received one readiness code per identified theme, and n = number of participants with that theme identified in transcript. The percentage of interview participants (B) coded as prepared in zero themes vs. one or more themes, by offer outcome (yes or no job offer). The denominator for each percentage is the total number of participants in that offer group. Readiness was coded from interview transcripts as Prepared, Mixed, Unprepared, or Unknown; only themes coded as Prepared are counted here.

We have pulled quotes from our interview transcripts with applicants. The following quote is from an applicant who received a job offer and perceived they were prepared because of mentorship from their PI. ‘My postdoc mentor has been a key figure in my preparation. He has been providing feedback on my application materials and helping me refine my research vision, and I’ve also consulted with senior faculties in my department and peers that have recently secured faculty positions.’ Another quote from an applicant who also received a job offer and remarked that their PI prepared them for the faculty job market: ‘My PI has a super excellent track record of placing people. She’s all about promoting her trainees and making sure she supports their careers. If I wasn’t ready to go on the job market, she would have flagged it. But she’s all in*.’*

Across all themes, ‘mixed’ ratings predominated over ‘prepared’, suggesting that participants more often demonstrated partial rather than complete self-reported readiness. Several themes — including personal context including family/geography and interdisciplinarity — had small coding bases (n = 10–11), limiting interpretation. The following is a quote from an applicant who received a job offer but had family constraints prior to going on the faculty job market: ‘This is my only shot. If this doesn’t work out, I’m out because of family restrictions. ‘Basically, we have these two small children — their ages are three and one — and we are financially strained because basically all of my salary is going to daycare right now.’

When we collapsed perceived readiness into a binary outcome, most participants in both groups had zero themes coded as Prepared: 65% of offer recipients (n = 20) and 75% of non-recipients (n = 12) (**Figure 4B**). A larger proportion of offer recipients had at least one theme coded as prepared compared to non-recipients (35% vs. 25%), though the small sample precluded formal statistical comparison.

**Figure 4.**
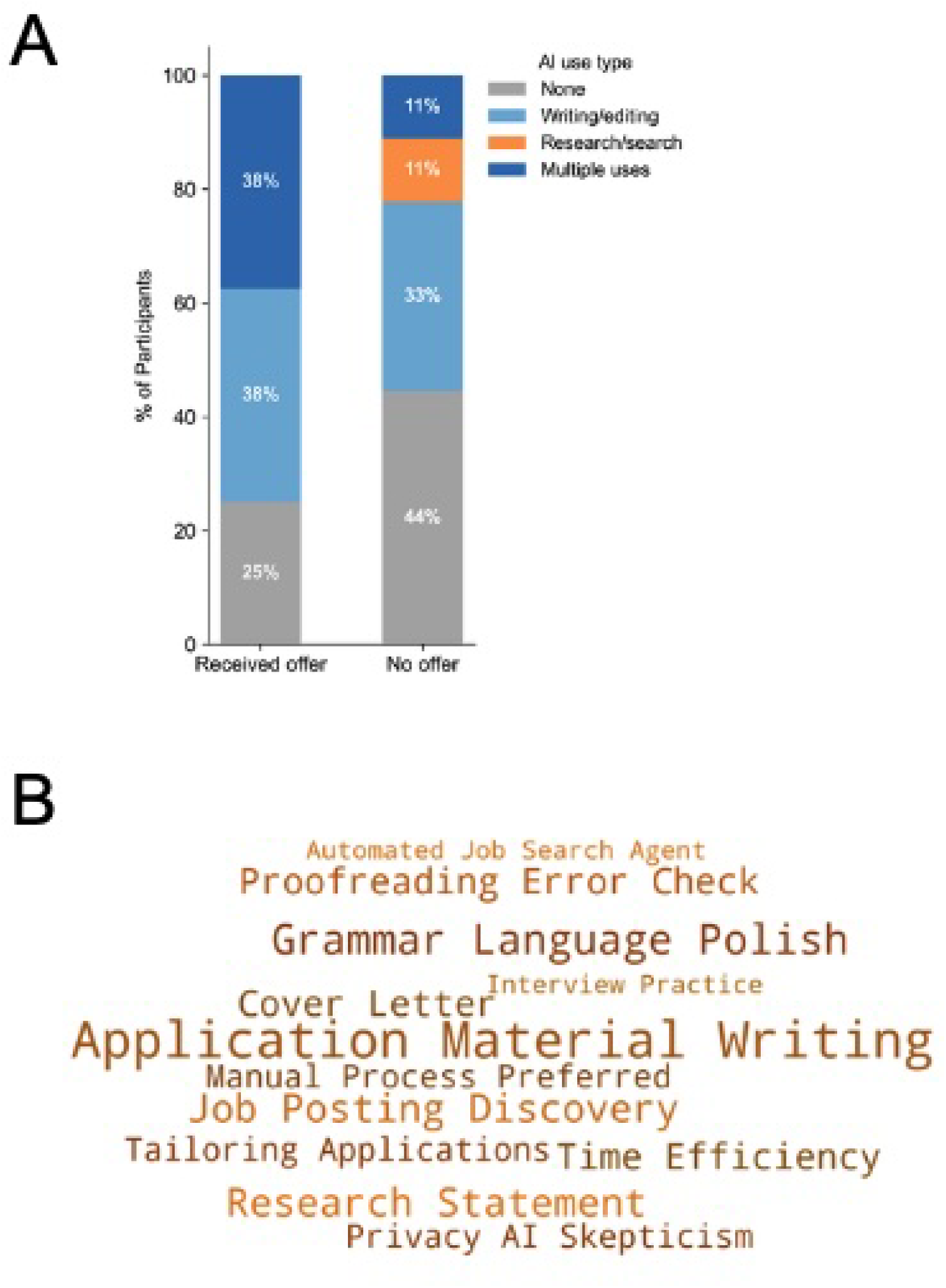
AI usage by applicants on the faculty job market. A, the percentage of participants that use artificial intelligence (AI) for faculty job market preparation. Categories of use include none (grey), writing and editing (light blue), research searches (orange) and multiple uses (dark blue). B, word cloud depicting what applicants used AI for while preparing for the faculty job market.

### Diverse usage of AI helps faculty job market applicants get an offer

With the emergent technology of generative AI appearing more mainstream during our data collection process, we had the unique opportunity to ask applicants about their use in their faculty job search. Among the 25 interview participants who were asked about AI tool use, those who received offers reported more diverse AI engagement than those who did not, with 38% reporting multiple use types compared to 11% of non-offer participants (**Figure 5A)**. Notably, research/search-only AI use was reported exclusively among non-offer participants (11%, n = 1), whereas offer recipients were more likely to use AI for writing and editing or for multiple purposes (**Figure 5B**). Though this difference did not reach statistical significance (Fisher’s exact p = 0.394), consistent with the limited power of this subgroup analysis, it suggests that part of being a ready applicant for the faculty job market may be an external signal of readiness.

**Figure 5.**
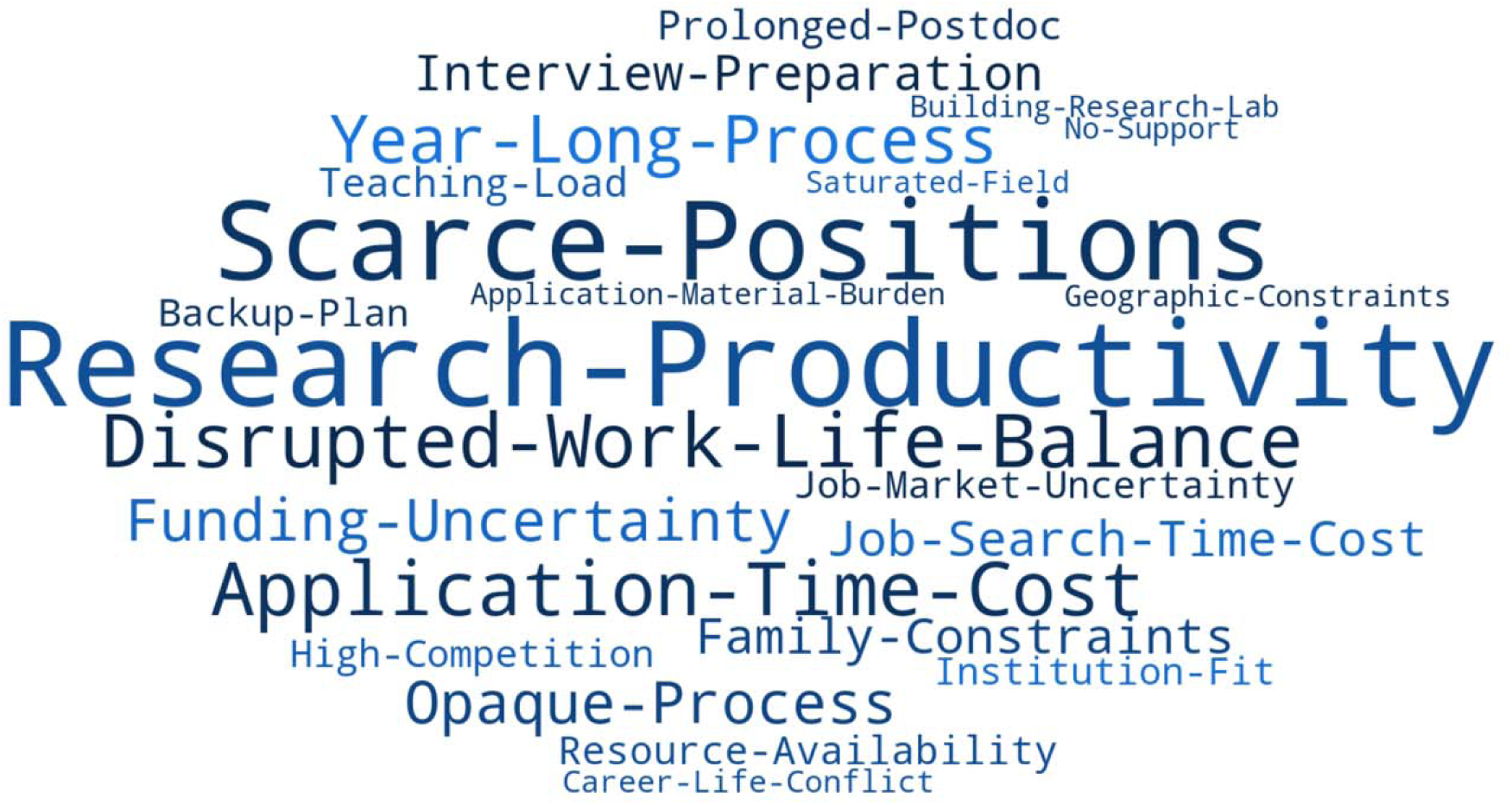
Word cloud depicting stressors reported by interviewed participants on the faculty job market. Phrases were constructed to reflect the specific nature of each stressor rather than its topic alone (e.g., Scarce-Positions rather than Positions). Word size is proportional to the number of participants who raised each theme; only themes endorsed by at least 4 of 29 participants (≥14%) are displayed. Each theme was counted at most once per participant regardless of frequency within their response.

### Being on the faculty job market is challenging in many areas for applicants

Our quantitative data suggests that there is a narrow window of time when applicants are competitive for a faculty job search, as discussed above and in our previous work^2,3^. Our qualitative data suggests that applicants can identify some themes that relate to their preparedness, but it is important to note that the faculty job market is unique in the fact that applicants must make this decision. Post-PhD positions are varied and nebulous but seen as necessary training. There are no determined “graduation” checkpoints that tell applicants when they are ready to enter the faculty job market. This is one example of many we have found throughout the years of doing this work of the lack of transparency “benchmark” and sources of stress around this important career transition^2^. In this new qualitative dataset, we asked applicants to describe sources of stress they associated with the academic faculty job market (**Figure 5**). Word cloud visualization of qualitative responses revealed that research productivity demands, and scarce positions were the most prominently endorsed themes, consistent with the quantitative readiness data showing research productivity as one of the most coded themes across participants. Application-time-cost and disrupted-work-life-balance emerged as prominent secondary stressors, reflecting the personal and logistical burden of the search process. Notably, opaque-process and year-long-process were frequently mentioned, suggesting that participants experienced the faculty market not only as competitive but as structurally non-transparent and prolonged. Family-constraints and geographic-constraints appeared as smaller but recurrent themes, highlighting that for many participants career decisions were inseparable from personal and relational circumstances.

## Discussion

A major challenge facing biomedical sciences trainees is determining when they are sufficiently prepared to enter the faculty job market. To address this gap, our group, the Faculty Job Market Collaboration (FJMC) has sought to provide evidence-based guidance for prospective applicants through a series of studies examining factors associated with faculty hiring success^1–4^. Building on this work, our recent research has focused specifically on the question of applicant readiness. We assembled a six-year longitudinal dataset from annual faculty job market surveys that captures key applicant characteristics and outcomes. Our analyses indicate that the strongest quantitative predictor of receiving a faculty job offer is the number of applications submitted and that there is an optimal window when you are competitive. To complement these findings, we conducted qualitative interviews with biomedical postdoctoral fellows who participated in the faculty job market. Among applicants who ultimately received offers, perceived readiness was strongly influenced by support from their principal investigator. However, feeling prepared in themes we identified as commonly viewed as indicators of readiness, including research productivity, grantsmanship, teaching and mentoring experience, networking, cohort support, personal and geographic considerations, mentorship, and interdisciplinarity, were not independently associated with obtaining a faculty position. We additionally observed that applicants who reported using AI tools across a broader range of job market activities experienced greater success. Collectively, our mixed-methods analyses demonstrate that faculty hiring outcomes were influenced by a complex interplay of factors, highlighting the need for data-driven approaches to assess applicant readiness and inform career decision-making.

The biomedical faculty job market experienced substantial disruption in 2024-2025 due to uncertainty surrounding U.S. federal research funding, particularly NIH support ^10,11^. Proposed reductions in indirect cost reimbursements, delays in grant review and funding decisions, and concerns about future federal research investments prompted many research universities to implement hiring freezes, delay faculty searches, reduce graduate admissions, and restrict spending^10,12^. These measures decreased the number of available faculty positions and intensified competition among applicants. At the same time, institutions and professional organizations warned that funding instability could weaken the biomedical research workforce pipeline by limiting opportunities for graduate students, postdoctoral researchers, and early-career investigators, creating one of the most challenging faculty hiring environments in recent years^12^.

The focus of our study was to determine at what point is a biomedical sciences postdoctoral fellow ready to go on the faculty job market. Our present work, combined with our previous studies^2,3^ strongly suggest that it is truly the number of applications an applicant submits per cycle that matters the most. This ideal data point has been reinforced by our 6-year LASSO regression model. From our qualitative interviews, it appears that PI/mentor/supervisor plays a role in helping applicants feel prepared to go on the job market. Among the 32 participants with known offer outcomes, the relationship between number of unprepared themes and likelihood of not receiving an offer was non-linear: participants reporting one unprepared theme had the lowest no-offer rate, while those reporting seven unprepared themes (n = 2) all received no offer. A U-shaped pattern in the data emerged that is intriguing but based on very small cell sizes. One of the two participants who reported seven unprepared themes is a K99 holder who did not receive an offer, this is a notable case. Our qualitative data sample size is small, and we are planning to build upon this data to determine whether we observe the same patterns in a larger pool of faculty applicants.

## Limitations

The objective of the present study was to identify factors associated with readiness for the biomedical faculty job market. Since 2018, we have collected annual survey data from applicants pursuing faculty positions in the biomedical sciences, resulting in a large longitudinal dataset that has enabled robust quantitative analyses of hiring outcomes and associated metrics. It should be noted that females are more likely to complete surveys^13^, we also see that in our data, ∼50.1% of respondents were females and 19.8% LGB+/GNC.

More recently, we expanded this effort to include a longitudinal qualitative study of biomedical postdoctoral scholars participating in a faculty job search cycle. Participants completed a minimum of three semi-structured interviews that focused on job market preparation, preparation for virtual interviews, and preparation for on-site interviews. Applicants who did not advance to subsequent stages of the search process completed a final interview addressing future career plans, whereas applicants who received faculty offers participated in an additional interview focused on negotiation and decision-making. This study generated a rich qualitative dataset capturing applicants’ experiences throughout the faculty hiring process. In the present analysis, we focused specifically on data obtained during the initial interview, which examined perceptions of readiness and preparation prior to entering the job market, and evaluated how these factors related to eventual hiring outcomes. Although the current sample is modest (n = 32), with 20 participants ultimately receiving at least one faculty job offer, these data provide an important preliminary foundation for understanding applicant readiness from the candidate perspective. Ongoing data collection across future hiring cycles will increase sample size and allow for more comprehensive analyses of the factors that contribute to success on the biomedical faculty job market. The integration of qualitative and quantitative approaches offers a unique opportunity to characterize the complex experiences of faculty applicants and develop evidence-based guidance for future trainees.

## Conclusions

Our mixed-methods analysis of the biomedical faculty job market indicates that there is no single criterion that determines applicant readiness or guarantees success. Rather, faculty hiring outcomes are influenced by a constellation of factors, including the breadth of applications submitted, the quality of mentorship and support from a principal investigator, strategic use of emerging tools such as artificial intelligence, and career stage characteristics. Notably, successful applicants often occupied senior postdoctoral or equivalent positions that demonstrated independence and leadership, while avoiding prolonged periods in training without clear evidence of progression. These findings underscore the complexity of the faculty hiring process, which involves multiple stakeholders and diverse evaluation criteria that vary across institutions and disciplines. By integrating quantitative and qualitative data, our evidence-based approach provides a more nuanced understanding of applicant readiness and offers practical guidance for trainees navigating the biomedical faculty job market.

## Supporting information

see supplementary file for survey questions

## Funding

These studies were funded by the Burroughs Wellcome Funding (Award Numbers: 1022092, 1417417; 1586063).

## Author contributions

Conceptualization: AH, YC, CTS, AJK, AKH, NMJ; Data curation: YC, AKH; Formal Analysis: YC; Funding Acquisition: AH, CTS, AJK, AKH, NMJ; Investigation: AH, YC, CTS, AJK, AKH, NMJ; Methodology: AH, YC, CTS, AJK, AKH, NMJ; Project administration: AH, YC, CTS, AJK, AKH, NMJ; Resources: AH, YC, CTS, AJK, AKH, NMJ; Software: YC, AKH; Supervision: AH, YC, CTS, AJK, AKH, NMJ; Validation: AH, YC, CTS, AJK, AKH, NMJ; Visualization: AH, YC, CTS, AJK, AKH, NMJ; Writing-original draft: YC, NMJ, AKH; Writing – reviewing and editing: AH, YC, CTS, AJK, AKH, NMJ

## Conflicts of interest

None declared.

## Disclaimers

None declared.

## Previous presentations

We have presented part of this work at the 2026 Annual National Postdoctoral Association meeting, March 13 to 14 2026, San Francisco, California.

## Data availability

All quantitative data is available on the Faculty Job Market Collaboration data dashboard https://www.faculty-job-market-collab.org

Qualitative interview data associated with this study cannot be made publicly available or shared upon request due to ethical constraints regarding participant privacy.

## Notes

https://www.faculty-job-market-collab.org/

### Competing Interest Statement

The authors have declared no competing interest.

