## Supplementary material for "When are Biomedical Postdocs Ready for the Faculty Job Market? A Mixed-Methods Analysis of Metrics and Resilience Among Faculty Job Seekers": see supplementary file for survey questions

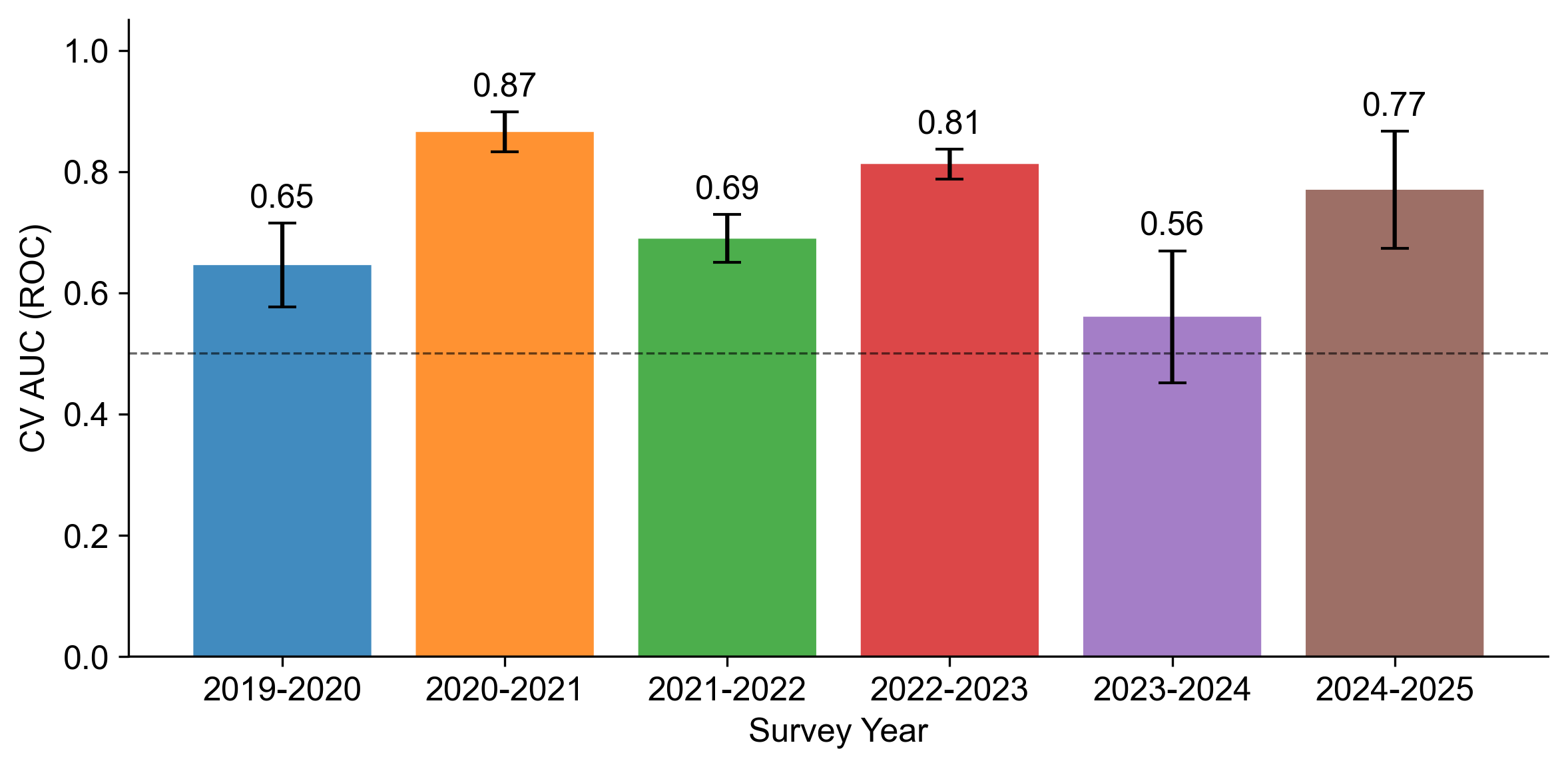

**Figure 1**. Cross-validated AUC by cycle. Dashed line = chance (0.50). Error bars = ± 1 SD across folds.

Supplementary materials for Haage A, Cheng Y, Smith C, Kozik AJ, Hagan AK, Jadavji NM^.^ When are Biomedical Postdocs Ready for the Faculty Job Market? A Mixed-Methods Analysis of Metrics and Resilience Among Biomedical Faculty Job Seekers

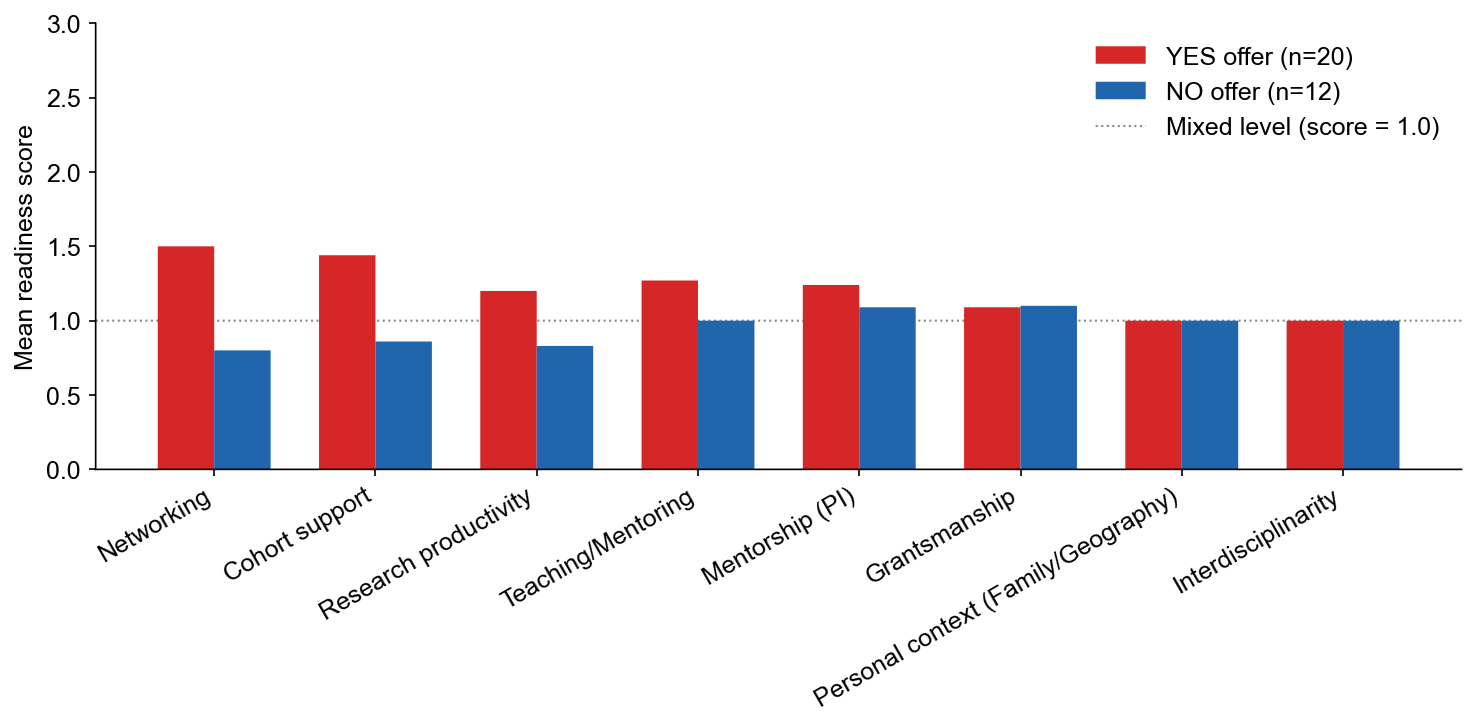

**Figure 2**. Theme-level associations between readiness and offer outcome were examined using three complementary approaches, each applied independently to each of the eight themes. (1) Fisher's Exact Test (2×2): compared the proportion coded Prepared between YES- and NO-offer groups, yielding an odds ratio (OR) and exact two-tailed p-value. (2) Exact Permutation Test (2×3): compared the full three-category readiness distribution (Prepared/Mixed/Unprepared) across offer groups using 5,000 permutations with the log-likelihood ratio statistic (G) as the test criterion; this extends the binary Fisher's test by preserving all three readiness categories as distinct. (3) Mann-Whitney U Test: readiness was encoded ordinally (Prepared = 2, Mixed = 1, Unprepared = 0) and distributions were compared non-parametrically, preserving the ordering of categories. Participants coded Unknown were excluded from all tests. Within each test, p-values across the eight themes were adjusted for multiple comparisons using the Benjamini-Hochberg (BH) false discovery rate procedure (α = 0.05; Bonferroni reference threshold = 0.00625). Analyses were conducted in Python 3.10.8 (SciPy 1.9.3, pandas 2.3.3).

Supplementary materials for Haage A, Cheng Y, Smith C, Kozik AJ, Hagan AK, Jadavji NM^.^ When are Biomedical Postdocs Ready for the Faculty Job Market? A Mixed-Methods Analysis of Metrics and Resilience Among Biomedical Faculty Job Seekers

All questions asked of respondents who completed our faculty job market survey from 2029 to 2025.

Quantitative Survey

What is your current position?

Have you ever previously held a tenure-track faculty or tenure-track equivalent faculty appointment?
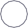
 Yes

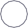
No

What is your gender?
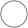
 Man

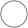
 Woman

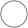
 Non-binary
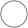
 Gender-fluid
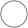
 Questioning

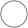
 Unlisted gender

Are you cis- or trans- identifying?
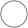
 Cis

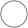
 Trans

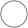
 Non-binary
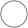
 Questioning

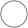
 Prefer not to say

What is your sexual orientation?
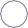
 Asexual

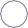
 Bisexual or bicurious

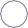
 Opposite-sex (heterosexual) only/mostly
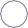
 Pansexual

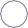
 Prefer not to say
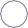
 Questioning

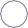
 Same-sex (homosexual) only/mostly

What is your race/ethnicity? Select all that may apply.
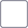
 Caucasian-American/European

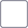
 Caucasian-American/North African or Middle Eastern
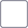
 African-American/Black/African

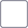
 Asian-American/Asian
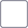
 South/Central American

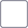
 North American Indigenous (e.g., Hawaii or Alaskan Native, American Indian)
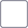
 North American Hispanic/Latinx

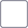
 Caribbean Islander (e.g., Puerto Rico, Trinidad)
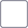
 Pacific Islander (excluding Hawaii)

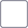
 Oceania (e.g., Australian/New Zealander)
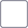
 Not Listed

Do you have a disability?

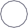
 Yes, I have a visible disability
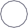
 Yes, I have a hidden disability
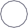
 No, I do not have a disability

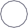
Choose not to respond

How old are you?

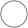
 20 - 25 years old

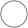
 26 - 30 years old

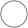
 31 - 35 years old

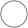
 36 - 40 years old

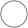
 41+ years old

You currently reside in	.
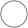
 The United States of America (U.S.)
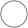
 Canada

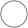
 Somewhere else

What is your legal status in the U.S. or Canada?
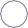
 Citizen

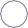
 Permanent resident

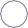
 Temporary work visa (e.g., H1B in U.S.)

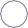
 Temporary student/scholar visa (e.g., F1, J1 in U.S.)
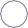
 Applying from outside the country(ies)

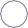
 Choose not to disclose

Are you a first generation (undergraduate) college graduate?
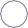
 Yes

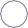
 No

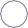
 Unsure

Are you a first generation Ph.D.?
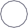
 Yes

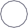
 No

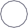
 Unsure

What is your relationship status?
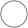
 Single

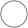
 Committed partnership(s)
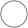
 Married

Do you have dependents (children/parents) in your household?
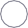
 No dependents

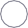
 Yes, one child

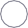
 Yes, multiple children

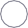
 Yes, adult(s) I/we take care of

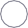
 Yes, adult(s) and child(ren) I/we take care of

Which category best fits your field of research?
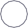
 Biological Sciences

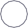
 Chemistry

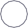
 Computer & Information Sciences
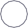
 Engineering

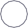
 Social, Behavior, & Economic Sciences
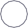
 Mathematical & Physical Sciences

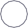
Geosciences

How old are you?

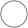
 20 - 25 years old

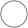
 26 - 30 years old

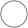
 31 - 35 years old

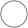
 36 - 40 years old

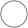
 41+ years old

You currently reside in	.
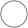
 The United States of America (U.S.)
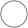
 Canada

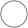
 Somewhere else

What is your legal status in the U.S. or Canada?
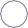
 Citizen

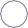
 Permanent resident

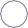
 Temporary work visa (e.g., H1B in U.S.)

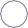
 Temporary student/scholar visa (e.g., F1, J1 in U.S.)
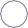
 Applying from outside the country(ies)

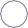
 Choose not to disclose

Are you a first generation (undergraduate) college graduate?
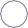
 Yes

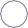
 No

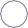
 Unsure

Are you a first generation Ph.D.?
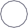
 Yes

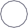
 No

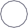
 Unsure

What is your relationship status?
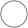
 Single

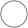
 Committed partnership(s)
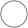
 Married

Do you have dependents (children/parents) in your household?
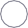
 No dependents

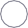
 Yes, one child

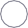
 Yes, multiple children

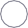
 Yes, adult(s) I/we take care of

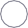
 Yes, adult(s) and child(ren) I/we take care of

Which category best fits your field of research? Biological Sciences

Chemistry

Computer & Information Sciences Engineering

Social, Behavior, & Economic Sciences Mathematical & Physical Sciences

Geosciences

Integrated Sciences Humanities

Does your research fall under Biomedical Sciences or Biomedical Engineering (e.g., involving public health or healthcare)? Yes

No

Unsure

Where did you complete your PhD? US

Canada

Internationally

**Core Metrics**

#### How many peer-reviewed papers have you been an author on?

#### If conference abstracts count towards publications in your field, how many of those have you been an author on?

#### Of peer-reviewed publications, how many papers have you published as corresponding author?

Of peer-reviewed publications, how many first author publications do you have?

How many pre-print papers have you posted online? If zero, please put "0"

How many *Cell Nature*, or *Science* publications are you an author on, TOTAL? If zero, please put "0"

How many first-author *Cell Nature*, or *Science* publications do you have? (If none, enter 0).

#### If you have a Google Scholar account, what is your number of citations (all)?

#### If you have a Google Scholar account, what is your number of citations (since 2020)?

#### f you have a Google Scholar account, what is your h-index (all)?

#### f you have a Google Scholar account, what is your h-index (since 2020)?

How many patents do you have approved or pending? 0

2

3

4

5

>5

Have you ever been awarded any of the following grants (select all that apply):

Predoctoral Fellowship Postdoctoral Fellowship

Transition to Independence Award (e.g., NIH K99) Research Project Grant (e.g., NIH R01), as co-PI Research Project Grant (e.g., NIH R01) as PI

Do you have any teaching experience? No

Yes, Teaching Assistant (TA) position Yes, experience beyond TA

If "Yes, experience beyond TA", please select all that apply:

Instructed undergraduate courses

Co-instructed undergraduate courses Guest lectured undergraduate courses Instructed graduate courses

Co-instructed graduate courses Guest lectured graduate courses

Independent instructor/lecturer (type of course not specified) Co-instructor/lecturer (type of course not specified)

Guest-lecture (type of course not specified) Lab course instructor

Lecturing for workshops

Instructor for high school courses Visiting assistant professorship

Adjunct teaching instructor for undergraduate courses at a community college or primarily undergraduate-serving institution

Adjunct teaching instructor for undergraduate courses at an R1, research-intensive university Other adjunct teaching positions

Teaching certificate

Teaching assistant for undergraduate or graduate courses

Does your CV list service projects, organizations, or committees?

Examples: Presenting to girl scout troups, leadership in your graduate student organization, or serving on the outreach committee of your field specfic society.....

Yes No

Please mark all service activities that appear on your CV. Member of graduate student organization

Leadership in graduate student orgnaization Membership in post-doctoral association

Leadership in post-doctoral association Founded a student club

Organized or led a seminar series Organized or led a journal club

Membership in a field specific society or organization Leadership in a field specific society or organization Community outreach

Founded an outreach organization Personal blog writing

Writing for a field specific society blog Peer review for journals in your field

Peer review for funding organizations in your field Membership in institution governance

Leadership in institution governance Tutoring

Mentoring outside of your research group

Quoted or contributed writing for popular press

Membership in an identity/affinity specific society or organization Leadership in an identity/affinity specific society or organization Union membership

Voting or non-voting participation on departmental search committee

Voting or non-voting participation on institutional search committee (ie, dean or leadership search)

How many service organizations, projects, or committees are listed on your CV? Please enter a numeric integer.

In what year did the **earliest** listed service organization/project/committee begin (e.g., 2019) ?

**Core Outcomes**

#### How many times have you applied for tenure-track/tenure-track equivalent positions?

If the 2024 - 2025 application cycle was the first time, please select "1", if you also applied last cycle, select "2", etc.

2

3

4

5

>5

### From May 2024 to April 2025, how many faculty job applications did you submit (TOTAL)? Use integer values (e.g., 1, 2)

### From May 2024 to April 2025, how many faculty job applications did you submit to research-intensive (R1 in US, U15 in Canada) institutions?

### From May 2024 to April 2025, how many faculty job applications did you submit to primarily undergraduate serving institutions?

From May 2024 to April 2025, how many **off site/early or initial round** interviews did you have TOTAL?

### Please list the full name of all institutions that contacted you for off site interviews, separated by semi-colons (Univ of A; X State Uni). Include the campus where appropriate. Do not assume that an acronym or shortened name will be sufficient (e.g. Columbia is both a college and a university).

These data will only be used for institutional classification purposes (e.g., Carnegie) and will not be shared outside of the research team. Please remember that all questions are optional.

From May 2024 to April 2025, how many **on-campus/later or final round** interviews did you have TOTAL?

### Please list the full name of all institutions that contacted you for on-campus/later or final round interviews, separated by semi-colons (Univ of A; X State Uni). Include the campus where appropriate. Do not assume that an acronym or shortened name will be sufficient (e.g. Columbia is both a college and a university).

These data will only be used for institutional classification purposes (e.g., Carnegie) and will not be shared outside of the research team. Please remember that all questions are optional.

During the May 2024 to April 2025 application cycle, how many faculty job offers did you receive TOTAL?

### Please list the full name of all institutions that gave you a formal written faculty job offer, separated by semi-colons (Univ of A; X State Uni). Include the campus where appropriate. Do not assume that an acronym or shortened name will be sufficient (e.g. Columbia is both a college and a university).

These data will only be used for institutional classification purposes (e.g., Carnegie) and will not be shared outside of the research team. Please remember that all questions are optional.

How did you respond to any offers received? Please check all that apply. I did not receive any offers

I accepted an offer at a primarily undergraduate institution I accepted an offer at a research-intensive institution

I rejected offer(s) for geographic fit

I rejected offer(s) for hostile/unwelcome interviewing environments I rejected offer(s) because the position did not fit my career goals

I rejected offer(s) because the personal compensation/benefits were insufficient I rejected offer(s) because the start up package was insufficient

I rejected offer(s) because of spouse/partner job prospects I rejected offer(s) because of other family needs

I rejected offer(s) for reasons not listed above

I am holding offer(s) until I can do a full in-person visit I am waiting on institutional approval of offer(s)

I rejected offer(s) for the political climate of the location

I rejected offers(s) because of the uncertainty of future federal funding

How many formal rejections did you receive (not including a lack of correspondence) TOTAL?

#### After participating in the academic job market this year (2024-2025), my commitment to an academic career has:

Become stronger Not changed

Become weaker

What are your plans for the next year if you **do not** accept a tenure-track faculty position by the conclusion of the 2024 - 2025 application cycle?

Have you withdrawn your application from any searches after doing either an initial zoom interview or an in-person interview? Yes - Please expand on why here

No

#### Do you want to comment on or emphasize something else about your experience on the academic job market that is not already represented in the questions above?

If you received a job offer(s) what was your salary?

If you received a job offer(s) what was your start-up amount?

What resources did you use to to prepare for the academic job market

Resources from Institution - Postdoctoral Office, Career Services, etc. Online tools

Future PI Slack

Other - Please specify

**Willing to Answer Further Questions?**

Would you be willing to answer additional questions regarding your **personal financial situation** for us to better understand how this intersects with job market success?

Examples include household income, student loan status, childcare, etc.

Yes

No, I don' t have time

No, I have privacy concerns

Would you be willing to answer additional questions on your **training environment**, including metrics of your PhD and/or post-doctoral advisor for us to better understand how this intersects with job market success?

Examples include resource used (e.g., Future PI slack), advisor H-index, etc. Yes

No, I don't have time

No, I have privacy concerns

Would you be willing to answer additional questions on your **perceptions and experiences of the interview process**?

Examples include intention to apply to faculty positions moving forward, unexpected interview questions/components, etc. Yes

No, I don't have time

No, I have privacy concerns

**Perceptions**

Please rate your level of agreement with the following statements.

Strongly

Disagree	Disagree	Neutral	Agree	Strongly Agree	N/A

I was adequately prepared for participating in the academic job market

My current supervisor was helpful and accommodating during this process

I am satisfied with the research products (materials, data, topics) my PI and I have agreed I will take with me to my new position

The processes involved in participating in the academic job market are transparent

I am satisfied with the outcome of my academic job search

My relationship status and/or family situation impeded my academic job search

My citizenship status impeded my academic job search

My mental health has been negatively impacted by my academic job search

I have not been able to complete other professional or personal goals directly due to time and energy spent on my academic job search

How were the finances for your travel to interviews handled? Were you ever asked to cover any costs upfront?

To what extent was your **service** discussed during the interview process? Never

Occasionally Frequently

Was your commitment to your research program ever questioned in light of your **service** activities?

Yes

No

To what extent was your **teaching experience** discussed during the interview process? Never

Occasionally Frequently

Was your commitment to your research program ever questioned in light of your **teaching activities**? Yes

No

During the interview process, did any **faculty member** ask about your marital and/or family status? No, never

Yes, only during remote interviews Yes, only during on-site interviews

Yes, during both remote and on-site interviews

During the interview process, did any **faculty member** ask about or question your willingness to relocate or otherwise accept the position?

No, never

Yes, only during remote interviews Yes, only during on-site interviews

Yes, during both remote and on-site interviews

During the interview process, did you experience any derogatory or otherwise offensive questions or comments regarding **your own**

Race/ethnicity

Nationality/Immigration Status Disability

Sexual orientation Gender identity

Religious observation (or lack thereof) Health status (including pregnancy)

During the interview process, did you experience any derogatory or otherwise offensive questions or comments regarding **someone else's**

Race/ethnicity

Nationality/Immigration Status Disability

Sexual orientation Gender identity

Religious observation (or lack thereof) Health status (including pregnancy)

Do you have additional comments regarding your **off-site/remote interview** experience(s)?

Do you have additional comments regarding your **on-site interview** experience(s)?

**Training Environment**

#### I solicited feedback on my application materials (CV, coverletter, research plan, etc.).

Yes No

#### This feedback came from (check all that apply):

My PI/advisor Peers

Family/Friends

Other faculty mentors (post-tenure) Other faculty mentors (pre-tenure)

Individuals in admin roles (dean, etc.)

#### I solicited feedback on my interview materials (chalk talk, seminar, teaching demo, etc.).

Yes No

#### This feedback came from (check all that apply):

My PI/advisor Peers

Family/Friends

Other faculty mentors (post-tenure) Other faculty mentors (pre-tenure)

Individuals in admin roles (dean, etc.)

#### I have an independent website or blog to host my CV information.

Yes No

#### I use social media at least in part to maintain a professional presence.

Yes No

#### I have attended workshops aimed at academic job market prep from my home institution.

Yes No

#### I am active in the peer-mentoring Future PI Slack group

Yes No

#### I have attended workshops aimed at academic job market prep from third party sources.

Yes

No

Please provide the academic job market prep workshop title and sponsor, if you would like:

I have used an Individual Development Plan (IDP) to aid in my decision to pursue a faculty career. Yes, and it was required by my program/training

Yes, but it was not a requirement of my program/training No

I found the IDP process helpful. Strongly agree

Agree

Somewhat agree

Neither agree nor disagree Somewhat disagree

Disagree

Strongly disagree

What is the faculty rank of your current advisor if you have one? Assistant Professor

Associate Professor Full Professor

From Google Scholar, what is the h-index of your current advisor?

From what institution did you receive your Ph.D. degree? Please provide the full institution name and include the campus where appropriate.

How many postdoc positions have you held (counting your current one if you have one)?

2

3

>3

List all institutions where you have served as a postdoc from first to last (most recent).

Please use full institution names and separate them with semi-colons (Univ of A; X State Uni) Include the campus where appropriate. Do not assume that an acronym or shortened name will be sufficient (e.g. Columbia is both a college and a university).

Number of times you personally reached out to faculty contacts at prospective institutions to ask questions about the department, institution, and support for new faculty. (numeric integer: 1, 3, etc...)

Check all that apply. For those institutions where I ultimately received an "onsite" interview, I first learned about the position(s) via: My postdoc advisor

My PhD advisor Job boards

Social media

A contact at the hiring institution

From an internal referral (the position was not advertised when I first learned about it)

Number of institutions **where I received an offer** AND I previously interacted with (met at conference, spoke via phone) at least one individual who was on the search committee BEFORE applying for the position. (Please use an integer response, e.g., 1, 2).

For the following questions, of the institutions where you had **on-campus interviews"**, we want you to indicate the number of institutions where this statement applies. Your answer should be an integer (1, 2, 3).

Someone on the search committee knows your PhD advisor.

Someone on the search committee has collaborated with your PhD advisor.

Someone on the search committee knows your most recent postdoc advisor.

Someone on the search committee has collaborated with your most recent postdoc advisor.

My PhD and/or postdoc advisor personally reached out to a contact at the institution to recommend me for the position.

How many conferences did you attend while "on the job market" during this cycle?

Did you speak to members of search committees from institutions where you were applying while at conferences during this cycle? Yes

No

I don't know

I believe networking and/or connections ultimately helped me obtain an "onsite" interview. Strongly agree

Somewhat agree Neutral

Somewhat disagree Strongly disagree

Check all that apply. For those institutions where I ultimately received an offer, I learned about the position(s) via:

My postdoc advisor My PhD advisor

Job boards Social media

A contact at the hiring institution

From an internal referral (the position was not advertised when I first learned about it)

**Finances**

What is your spouse/partner's occupation? I do not have a spouse/partner

Postdoc

Pre-tenure or research/teaching faculty Tenured/Tenure-track faculty

Employed (not academia or higher education)

Employed (elsewhere in academia/higher education) Self-Employed

Unemployed

Primary caregiver (stay at home parent) Student (undergraduate or graduate)

What is the highest level of education completed by your spouse/partner? I do not have a spouse/partner

Middle School (grade 8) High School (grade 12) Bachelors (4-year degree)

Associates (2-year degree) Vocational/Trade School Masters

Ph.D.

Professional Graduate Program (MD, JD, Physician assistant etc.)

What was your combined household income in US dollars (before taxes) last year (2024)? $25,000 - 30,000

$30,001 - 50,000

$50,001 - 70,000

$70,001 - 90,000

$90,001 - 120,000

$120,001 - 150,000

More than $150,000

Are you currently paying student loans? Yes

No, they are in deferment

No, I used non-loan financial aid (e.g., grants, scholarships) No, I did not use financial aid

No, I have already paid them or received loan forgiveness

Do you have any income from rentals, interest, dividends, or capital gains? Yes

No

Do you have friends, family, or similar network that could provide financial support in case of an emergency? No

Yes, but only up to $500 Yes, but only up to $1000

Yes, in amounts greater than $1000

How many children do you have that are too young for school and require childcare (i.e., under age 6)?

Is the income from your current (or most recent) position sufficient for the cost of living in your area? Yes

No

If "no" to above, how do you manage the financial gap? (Check all that apply) Credit cards

Extremely frugal living (e.g., noodles/beans/rice for dinner every night) Depend on income from a spouse/partner

Require a roommate to afford housing

Depend on financial loans/gifts from family

Require supplemental income from part-time/freelance work

What percent of your total household income do you estimate that you spend on housing costs?

What percent of your total household income do you estimate that you spend on childcare costs?

Supplementary materials for Haage A, Cheng Y, Smith C, Kozik AJ, Hagan AK, Jadavji NM^.^ When are Biomedical Postdocs Ready for the Faculty Job Market? A Mixed-Methods Analysis of Metrics and Resilience Among Biomedical Faculty Job Seekers

**Table 1.** Demographics of biomedical sciences post-doctoral fellows from 2024-25 and 2025-26 qualitative data collection.

| Total Respondents | 35 |
| --- | --- |
| **Gender** |  |
| Female | 21 |
| Male | 14 |
| LGB + GNC | 0 |
| Prefer to no say | 1 |
| **Current Institution** |  |
| Very High Research Activity (R1) | 31 |
| High Research Activity (R2) | 3 |
| Master's colleges and universities: larger programs (M1) | 1 |
| Asian | 5 |
| Black/African American | 9 |
| Hispanic or Latinx | 4 |
| Middle Eastern | 1 |
| White | 16 |

Supplementary materials for Haage A, Cheng Y, Smith C, Kozik AJ, Hagan AK, Jadavji NM^.^ When are Biomedical Postdocs Ready for the Faculty Job Market? A Mixed-Methods Analysis of Metrics and Resilience Among Biomedical Faculty Job Seekers

**Questions asked during first interview with biomedical sciences postdoctoral fellows preparing for the faculty job market during the 2024-25 and 2025-26 hiring cycles.**

**INITIAL INTERVIEW**

**Getting Acquainted & Rapport Building**

1. If you could just start by telling us a little about your professional and educational background beginning with choosing your doctoral program.
2. Reflecting on selecting your doctoral program, what were the motivating factors in that decision?
   a. Was pursuing a faculty position your goal at that time? How, if at all, did that goal factor into your decision-making process?
      i. If not, when did you decide that you wanted to pursue a faculty career? What were the motivating factors in that decision?
   b. What other factors influenced your decision-making process?
      i. Personal Identities or Contexts (e.g., partner, children)?
      ii. Faculty mentors/Prestige?
      iii. Geography?
   c. Knowing what you know now, how satisfied are you with that decision and experience?

**Choosing & Transitioning to a doctoral degree**

3. I would like to learn about your selection of a doctoral degree institution
   a. Why did you choose the institute? Research area?
   b. Research area of focus
   c. Institutional type
   d. Research interests (for a lay person)
   e. Did you use any campus-based resources (e.g., career services), professional resources, or support programs?
4. Next, I’d like to learn about your postdoc and research interests.
   a. How many postdocs have you done?
   b. Did you continue in your PhD lab for a postdoc? Or did you start in a new lab for your postdoc?
   c. Research area of focus
   d. Institutional type and public/private
   e. Research interests (for a lay person)
5. Let’s talk about how you chose your postdoc. Can you talk us through your process for choosing your position?
   a. How did you learn about postdocs and the process for securing one?
   b. How many did you apply for?
   c. How did you hear about postdoc opportunities?
   d. What were the primary criteria you used for identifying positions? Which of those were most important in your final decision?
      i. How did your future career aspirations influence your postdoc institution election?
   e. How many accepted you? Offered funding? If you received funding how long was it for?
   f. What support systems did you use during the decision process?
      i. What role did mentors or others play in your decision-making process?
      ii. How, if at all, have your postdoc positions and/or supervisors influenced your search process?
      iii. Did you use any campus-based resources (e.g., career services), professional resources, or support programs?
           1. If yes, how did you choose the resources?
           2. How was your experience? What was most helpful? What, if anything, was not helpful?
   g. Are you part of a cohort model program (e.g., K99 awardee, BWF program)? If so, what was that experience like for you?
      i. How did the cohort support you?
      ii. What worked well?
      iii. What could have been improved?
   h. Do you consider your research as interdisciplinary in nature? If so, did this influence the choice of your department or institution?
   i. Knowing what you know now, how satisfied are you with your postdoc decisions and experiences?
   j. How has your postdoc experience(s) influenced your desire, approach, or experience pursuing a faculty position?

**Faculty Interest**

Now, I want to learn about your interests in a faculty position and things that are influencing your job search process.

6. If you could write your own position description and work at any institution you wanted, what would be your ideal position and institution?
7. What types of positions do you anticipate applying for?
   a. Does the nature of your research (e.g., being interdisciplinary) influence the choice of positions or institutions?
8. What are the current factors that will guide your decision about applying for positions?
9. How many positions do you anticipate applying for?
10. How prepared do you currently feel to enter the faculty job market?
    a. What strategies and approaches have you taken to prepare?
       i. When did you begin preparing for the job market?
       ii. How has personal performance (e.g., previous publications, conference presentations, grants) contributed to your sense of preparation?
       iii. What roles have others played in your preparation?
    b. What role, if any, has AI played in your job search process and preparation?
11. What stresses you about entering the faculty job market?
12. Are there any aspects that you feel impact how you approach this search?
    a. Personal Identities (e.g., race, gender, disability) or Contexts (e.g., partner, children)?
    b. Faculty mentors/Prestige?
    c. Geography? (e.g., proximity, politics)
13. What else is important for us to know that we have not asked about? Is there anything you thought we might talk about that we haven’t yet?

Can keep this congruent with the beginning question (more explanation is provided here, for example, for personal identities, that is not provided above)

Perhaps also consider asking about Interdisciplinary nature of their work and choice of department. Also, consider asking in general about mentors and their role in this process similar to some of my comments above…essentially, ensuring to ask about mentor and career resources at each of the three topic areas of these questions (doctoral choice, postdoc choice, faculty choice) so those could be compared…and make sure there aren’t other factors you’ve asked about at each stage that are left out so they can all be compared across the stages. Somewhere across these questions, asking about cohort models (availability of those, or their participation in these), since they are known, especially, to support students from underrepresented backgrounds in continuing to pursue science/fac

**NEXT STEPS**

- You must complete a brief form after each interview to verify participation. Please use this link to verify the completion of today’s interview: <https://forms.office.com/r/f4p8NTJEAe>
- At the completion of the entire study, you will receive a gift card with compensation for your participation in the study ($75/interview completed).
- Follow up in by the end of February for interview #2
